# *miR-210* controls the evening phase of circadian locomotor rhythms through repression of *Fasciclin 2*

**DOI:** 10.1101/399873

**Authors:** Wesley Leigh, Zhenxing Liu, Xiaoge Nian, Yong Zhang

**Affiliations:** Department of Biology, University of Nevada Reno, Reno, NV 89557, USA; Department of Entomology and MOA Key Lab of Pest Monitoring and Green Management, College of Plant Protection, China Agricultural University, Beijing, 100193, China.

## Abstract

Circadian clocks control the timing of animal behavior rhythms to anticipate daily environmental changes. Fruit flies gradually increase their activity and reach a peak of activity around dawn and dusk. microRNAs are small non-coding RNAs that play important roles in post-transcriptional regulation. Here we identify *Drosophila miR-210* as a critical regulator of circadian rhythms. Under light-dark conditions, flies lacking *miR-210* (*miR-210*^*KO*^) exhibit a dramatic phase advance of evening anticipatory behavior about 2 hours. However, circadian rhythms and molecular pacemaker function are intact in *miR-210*^*KO*^ flies under constant darkness. Furthermore, we identify that *miR-210* determines the evening phase of activity through repression of the cell adhesion molecule *Fasciclin 2 (Fas2).* Ablation of the *miR-210* binding site within the 3’ UTR of *Fas2* (*Fas2*^*ΔmiR-210*^) by CRISPR-Cas9 advances the evening phase as in *miR-210*^*KO*^. Indeed, *miR-210* genetically interacts with *Fas2*. Moreover, Fas2 abundance is significantly increased in the optic lobe of *miR-210*^*KO*^ and *Fas2*^*ΔmiR-210*^. In addition, overexpression of *Fas2* in the *miR-210* expressing cells recapitulates the phase advance behavior phenotype of *miR-210*^*KO*^. Together, these results reveal a novel mechanism by which *miR-210* regulates circadian locomotor behavior.

**Author summary:** Circadian clocks control the timing of animal physiology. *Drosophila* has been a powerful model in understanding the mechanisms of circadian regulation. Fruit flies anticipate daily environmental changes and exhibit two peaks of locomotor activity around dawn and dusk. Here we identify *miR-210* as a critical regulator of evening anticipatory behavior. Depletion of *miR-210* in flies advances evening anticipation. Futhermore, we identify the cell adhesion molecular *Fas2* as *miR-210’s* target in circadian regulation. Fas2 abundance is increased in fly brain lacking of *miR-210*. Using CRISPR-Cas9 genome editing method, we deleted the *miR-210* binding site on 3’ untranslated region of *Fas2* and observed similar phenotype as *miR-210* mutants. Altogether, our results indicate a novel mechanism of *miR-210* in regulation of circadian anticipatory behavior through inhibition of *Fas2*.

## Introduction

The circadian locomotor rhythms of flies are generated by a neuronal network, which consists of ∼150 brain circadian neurons expressing core pacemaker genes [1-4]. These neurons can be further divided into 7 clusters based on their cell localization and neurotransmitters they express [4]. There are 3 groups of dorsal neurons (DN1, DN2, and DN3), 2 groups of ventral-lateral neurons (large and small LNv), dorsal lateral neurons (LNd), and lateral posterior neurons (LPN) [4]. The large LNvs (lLNvs) and 4 small LNvs (sLNvs) express the neuropeptide Pigment Dispersing Factor (PDF), while the 5^th^ sLNv is PDF negative. Under regular light-dark (LD) conditions, flies exhibit a bimodal pattern of locomotor rhythms, peaking around dawn and dusk, which are termed morning peak and evening peak, respectively. According to the dual-oscillator model, separate morning and evening oscillator track these behavior peaks separately [5]. Indeed, studies have shown that the PDF-positive sLNvs are responsible for promoting the morning peak, while the LNds, as well as the fifth sLNv, are mainly responsible for generating the evening activity [6, 7]. In addition, the DN1s are important to integrate environmental inputs such as light and temperature, and regulate circadian rhythms and sleep behavior [8, 9]. A recent study indicated that DN1s sense acute temperature fluctuations to modulate fly sleep [10].

The PDF positive sLNvs are the master pacemaker neurons in fly brain: they synchronize other circadian neurons to maintain robust circadian rhythms under constant darkness [11]. These sLNvs send axonal projections toward the dorsal brain region, where the DN1s and DN2s are located [12, 13]. The dorsal projections of sLNvs exhibit circadian arborization rhythms with higher complexity of axon terminals found in the early day and lower complexity at the night [14]. The physiology relevance and molecular mechanisms underlying this structural plasticity of sLNvs remain unclear. Recently, the cell adhesion molecule *Fasciclin 2* (*Fas2*) has been found to regulate the arborization rhythms of sLNvs [15]. In addition, two matrix metalloproteinases, MMP1 and MMP2 as well as the pacemaker protein Vrille were shown to be required for the structural remodeling control of sLNv projections [16, 17].

Circadian rhythms are generated by an intracellular molecular clock, which is conserved across the animal kingdom [18]. The core of this molecular pacemaker is a negative transcriptional-translational feedback loop [19-21]. In *Drosophila*, CLOCK (CLK) dimerizes with CYCLE (CYC) to activate rhythmic transcription of hundreds of genes through the E-box region [22, 23]. Among these clock-controlled genes, PERIOD (PER) and TIMELESS (TIM) are the key transcription repressor: they form heterodimers in the cytoplasm, and then enter into the nucleus to block their own transcription. The abundance of PER and TIM is tightly regulated by a series of post-translational modifications such as phosphorylation, glycosylation and ubiquitination [24-27]. Recent research has revealed an abundance of post-transcriptional regulation of circadian rhythms by both RNA binding proteins and miRNAs [28-30].

miRNAs are small non-coding RNAs, which repress target gene expression through mRNA degradation and/or translation inhibition, thus play crucial roles in post-transcriptional regulation [31]. Recent studies have uncovered functions of the miRNA biogenesis pathway and specific miRNAs in the regulation of different aspects of animal circadian rhythms [30, 32-34]. For instance, mouse *miR-132/212* modulates the seasonal adaptation and circadian entrainment to day length [35]. miRNAs also target the molecular clock to control period length or rhythmicity of circadian locomotor activity [36]. In *Drosophila*, the miRNA *bantam* regulates circadian locomotor period by repressing *clk*, while *miR-276a* and *let-7* inhibit the clock genes *timeless* and *clockwork orange* to modulate circadian rhythms respectively [32, 37, 38]. Moreover, various circadian output pathways are controlled by miRNAs. The *miR959-964* cluster miRNAs regulate the circadian timing of feeding and immune response, while *miR-279* modulates circadian locomotor behavior output [39, 40]. Recently, we and others have demonstrated that *miR-124* specifically controls the phase of circadian locomotor rhythms under constant darkness [41, 42].

Previously we identified that depletion of GW182, the key protein for miRNA function, affects PDF-receptor signaling and disrupts circadian rhythms [34]. To further understand the roles of miRNA, we performed a genome-wide screen for miRNAs controlling circadian rhythm phenotypes. Here we show that a conserved miRNA, *miR-210*, regulates circadian rhythms in *Drosophila*. *miR-210* determines the phase of evening anticipatory behavior through inhibition of *Fas2*.

## Results

### A genetic screen of miRNA mutants identifies *miR-210* as a regulator of circadian behavior

To identify miRNAs that regulate circadian rhythms, we screened the circadian locomotor rhythms of available miRNA mutants from Bloomington stock center. Circadian behavior of 46 lines in total was observed both under constant darkness (DD) and light dark (LD) cycles. Most of the miRNA mutants we tested exhibited normal circadian rhythms under DD; though, a few miRNAs had a slightly altered period or rhythmicity (Table S1). We then closely examined for the locomotor behavior under LD. Interestingly, we found that the *miR-210* mutant (*miR-210*^*KO*^) clearly advances the phase of the evening anticipatory peak about 2 hours (Figure 1A, 1B). The *miR-210*^*KO*^ was generated by knocking-in a GAL4 cDNA in replacement of the endogenous *miR-210* sequence [43]. To rule out the possibilities of off-target or genetic background effects, we utilized the *miR-210* knock-in mutant to perform a rescue. By restoring the expression of *miR-210*, we successfully corrected the evening anticipation in the *miR-210*^*KO*^ (Figure 1A, 1B).

**Figure 1:**
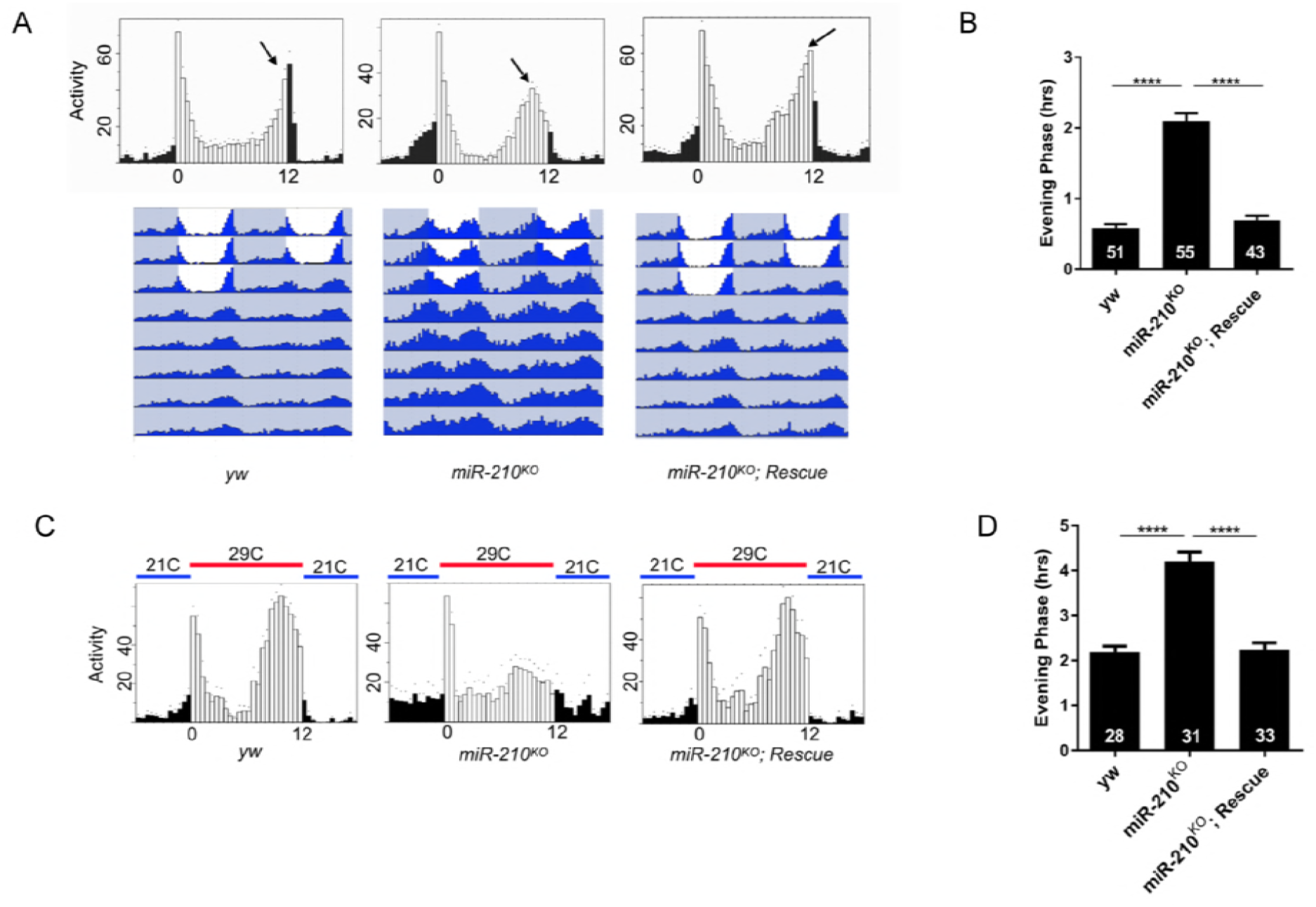
Loss of *miR-210* advances evening peak activity under LD and temperature cycles (TC). (A) Upper panel: representative eduction profiles of fly locomotor activity under 12:12 LD cycle. Arrow indicates the peak of evening activity. Black represents the dark phase, while white represents the light phase. Eduction is analyzed based on average of 3 days LD. Lower panel: representative double plotted actograms of flies under LD and DD conditions. Grey shadow represents the dark phase, while white represents the light phase. (B) Quantification of evening phase of flies under LD. Phase point is determined by the time differences between the peak of evening activity and light off time. Number of flies tested is listed in each bar. Error bars indicate Standard Error. ****= p<0.0001, determined by Student’s t test. (C) Representative eduction profiles of flies under 29°C: 21°C temperature cycles (TC) in DD. Blue represents 21°C for 12 hours, while red represents 29°C. (D) Quantification of evening phase of flies under TC. Error bars represent Standard Error. **** = p<0.0001 was determined by Student’s t test.

To examine whether the regulatory role of *miR-210* on evening anticipatory behavior is specific to light, we entrained the flies under 29°C: 21°C cycles (TC) in constant darkness. Under TC, the evening anticipation of wild-type flies exhibited an advance about 2.5 hours (Figure 1C, 1D) [44]. In *miR-210*^*KO*^, the overall activity was decreased under TC, however, we still observed an additional 2-hour phase advance compared to the controls (Figure 1C, 1D). Furthermore, we were able to rescue this behavior defect. These data confirmed that *miR-210* is a clear regulator of evening phase during entrainment.

The phase advance of evening anticipation in *miR-210*^*KO*^ is reminiscent of the *pdf* mutant phenotype [11]. To test the genetic interaction of *miR-210* and the PDF pathway, we generated *miR-210*^*KO*^; *pdf*^*01*^ double mutants. Under 12:12 LD conditions, the phase advance of the evening peak in *miR-210*^*KO*^ was indistinguishable from the *pdf*^*01*^ mutants (Figure S1A). No additive effect on the phase advance was observed in the *miR-210*^*KO*^; *pdf*^*01*^ double mutants comparing with *miR-210*^*KO*^ or *pdf*^*01*^ (Figure S1A, S1B). To observe the phenotype even more clearly, we tested these flies under long photoperiods with 16:8 LD cycle so that we avoid acute effects of the light-off transition on the shape of the evening peak. As observed in *pdf*^*01*^ mutant, *miR-210*^*KO*^ advanced the phase of evening anticipation by about 4 hours (Figure S1C, S1D). Again, no additive phase advance was observed in the *miR-210*^*KO*^; *pdf*^*01*^ double mutants (Figure S1D). These data suggest that *miR-210* functions in the same pathway as PDF in regulation of evening phase under entrainment.

To further test the role of *miR-210*, we first overexpressed it in all circadian tissues using *tim-GAL4* [45]. However, most of the flies with overexpression died rapidly after the LD cycles, which indicates that overexpression leads to severe health issues (Table 1). Thus, we then used the *pdf-GAL4* to specifically drive *miR-210* expression in PDF positive pacemaker neurons. The majority of flies (∼63%) with *miR-210* overexpression in PDF neurons became arrhythmic under DD. However, for the 37% rhythmic flies, the circadian period was lengthened about 2 hours. Thus, while loss of *miR-210* has no effect on the period and amplitude of circadian rhythms in DD, overexpression is detrimental to the proper function of the circadian oscillator in PDF neurons.

**Table 1:** Locomotor activity of flies with altered *miR-210* and *Fas2* in DD.

| Table 1: Locomotor activity of flies in DD |  |  |  |  |
| --- | --- | --- | --- | --- |
| Genotype | N | % Rhythmic | Period (hr) ± S.E.M. | Power<sup>#</sup> ± S.E.M |
| *tim-GAL4/+; Pdf-GAL80/+ M* | 42 | 100.0±0.0 | 24.5±0.4 | 94.4±2.6 |
| *tim-GAL4/+; Pdf-GAL80/*<br>*UAS-miR-210 M* | 35 | N/A* | N/A* | N/A* |
| *tim-GAL4/+ M* | 40 | 96±2.3 | 24.1±0.3 | 97.2±2.5 |
| *tim-GAL4/UAS-miR-210 M* | 33 | N/A* | N/A* | N/A* |
| *Pdf-GAL4/+ M* | 41 | 96.4±1.4 | 24.6±0.2 | 98.4±1.1 |
| *Pdf-Gal4/UAS-miR-210 M* | 44 | 37.1±3.5 | 26.1±0.6 | 65.0±1.5 |
| *yw M* | 45 | 92.3±2.3 | 24.7±0.2 | 91.3±1.8 |
| *yw F* | 42 | 91.6±3.6 | 24.1±0.4 | 88.7±2.1 |
| *miR-210<sup>KO</sup> M* | 52 | 88.0±2.4 | 24.2±0.4 | 87.9±3.2 |
| *miR-210<sup>KO</sup> F* | 49 | 87.5±3.1 | 24.5±0.3 | 89.6±1.6 |
| *miR-210<sup>KO</sup>/+ F* | 31 | 90.2±1.9 | 24.3±0.1 | 91.4±2.8 |
| *miR-210<sup>KO</sup>; UAS-miR-210 M* | 43 | 90.9±2.1 | 24.6±0.3 | 86.4±3.1 |
| *Fas2<sup>ΔmiR-210</sup> M* | 43 | 93.2±1.8 | 24.1±0.2 | 92.9±2.4 |
| *Fas2<sup>ΔmiR-210</sup> F* | 39 | 94.4±0.8 | 24.1±0.3 | 89.8±3.1 |
| *Fas2<sup>ΔmiR-210</sup>/+ F* | 43 | 95.3±1.7 | 24.3±0.1 | 93.8±4.2 |
| *miR-210<sup>KO</sup>/Fas2<sup>ΔmiR-210</sup> F* | 35 | 88.6±1.7 | 23.9±0.4 | 87.8±2.1 |

<sup>#</sup>Power is a measure of rhythm amplitude and corresponds to the height of the periodogram peak above the significance line.
\*N/A, flies were all dead after entering the constant darkness.

### The molecular pacemaker is intact in the *miR-210*^*KO*^ flies

Since circadian locomotor rhythms are unaffected in *miR-210*^*KO*^ flies under DD, it is likely that the molecular pacemaker in circadian neurons of *miR-210*^*KO*^ flies is functional. To confirm this, we examined the oscillation of the key pacemaker protein PER in DD. We first focused on the PDF positive sLNvs, which are the master pacemaker neurons under constant conditions. As expected, we found no obvious changes in PER oscillation or abundance in these sLNvs (Figure S2A, S2B). As an important output molecule for circadian rhythms, the PDF abundance was not affected in *miR-210*^*KO*^ flies (Figure S2C). We further examined PER levels in another two important groups of circadian neurons: LNds, and DN1s. Similar as in sLNvs, PER oscillation in these neurons were not affected (Figure S3). Taken together, these data suggest that loss of *miR-210* has no effects on the molecular clock.

### Loss of *miR-210* disrupts the circadian arborization rhythms of sLNv dorsal projections

Although genetic interactions between *miR-210* and the PDF pathway was identified (Figure S1), we observed no significant changes in PDF abundance in the flies missing *miR-210* (Figure S2C). Thus, we examined the projections of the sLNvs. Circadian arborization rhythms of the dorsal projections of sLNvs have been observed [14, 15]. To determine whether *miR-210* affects the arborization rhythms of sLNv projections, we used a PDF-specific antibody to examine the termini of sLNv dorsal projections in *miR-210*^*KO*^ flies at early day (Zeitgeber time 2 (ZT2), ZT0 is light on and ZT12 is light off) and early night (ZT14). We found that the wild-type flies had more PDF positive branches of axon terminals at ZT2 than at ZT14 (Figure 2A-2C), as previously observed [15]. Remarkably, this arborization rhythm was abolished in the *miR-210*^*KO*^ flies (Figure 2A-2C). This is mainly due to the significant decrease of axonal crosses at ZT2 compared to control flies (Figure 2A). Restoration of *miR-210* expression rescued the arborization rhythm in *miR-210*^*KO*^ mutants. Moreover, we did not detect significant changes of PDF abundance in the sLNv soma, indicating that the decrease of axonal branches was not due to the decrease of PDF staining (see Figure S2). Together, these results indicate that *miR-210* is required for the circadian arborization rhythms in the sLNv dorsal projections.

**Figure 2:**
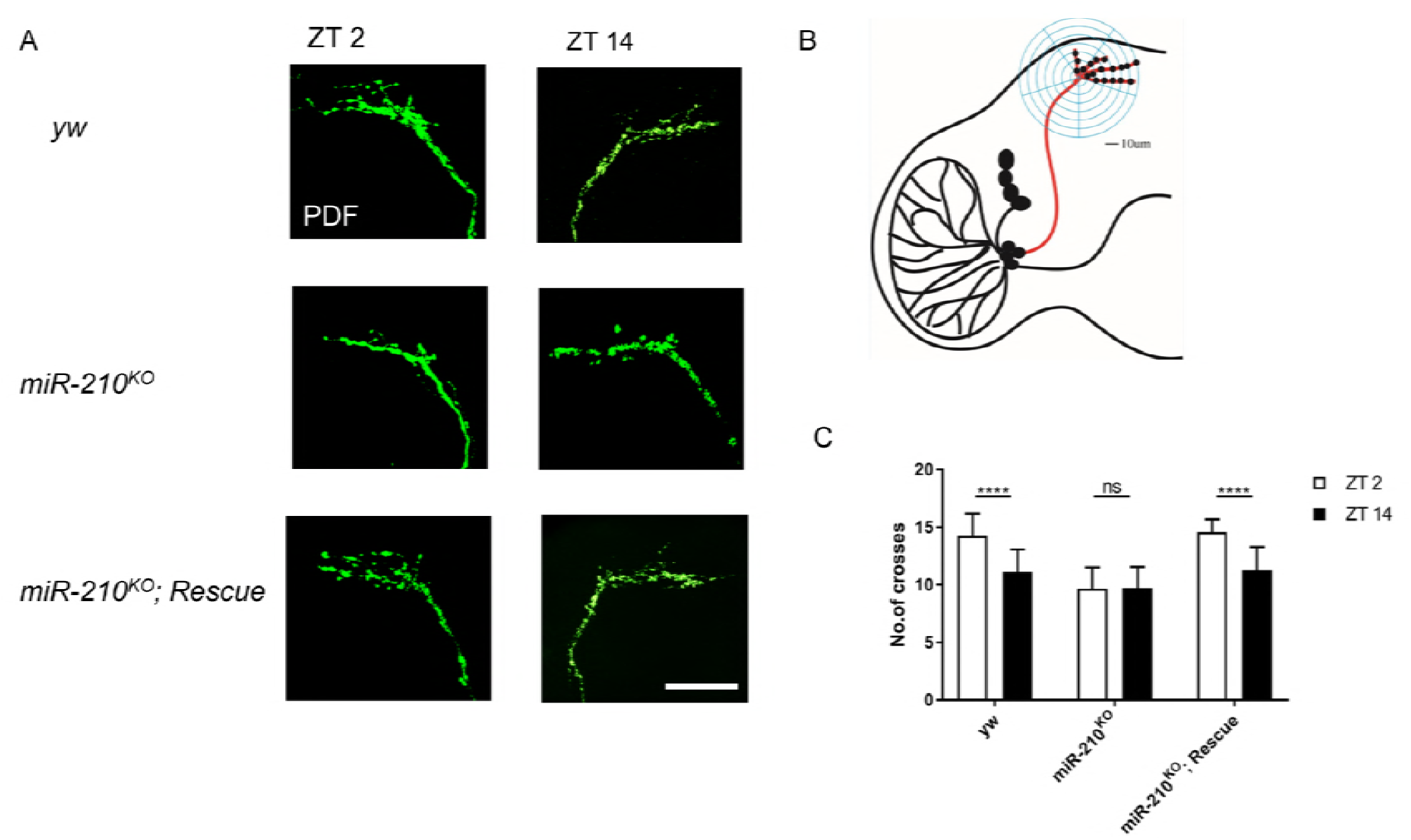
Loss of *miR-210* disrupts the circadian arborization rhythms of sLNv dorsal projections. (A) Representative confocal images of sLNv dorsal axonal projections. The brains were stained with anti-PDF antibody (green) at ZT2 at the 3rd day of LD. Scale bar is 65 μm. Schematic illustrating a modified Sholl’s analysis for quantification axonal termini defasciculation of sLNv neurons. See [15] and methods part for details. (C) Quantification of axonal morphology (fasciculation) of sLNv dorsal termini. The plot represents number of crosses between axonal branches and concentric rings. Plots show mean values, error bars indicate Standard error. **** represents p-value < 0.0001 by Student’s t test. ns represents not significant.

### *miR-210* controls the phase of evening peak by inhibition Fas2

miRNAs play negative posttranscriptional functions through binding to the 3’ untranslated regions (UTRs) of target mRNAs [31]. We searched putative *miR-210* targets by using an *in silico* prediction algorithm (http://www.targetscan.org/fly_12/), and identified *Fas*2 as one of the potential targets (Figure 3A). The putative *miR-210* binding sites within the 3’UTR of *Fas2* are highly conserved across *Drosophila* species (Figure 3A). Interestingly, Sivachenko et al. have reported that overexpression of *Fas2* in the PDF neurons caused fasciculation of sLNv dorsal projections and disrupted arborization rhythms [15]. Since miRNAs are normally negative regulators of target genes, and overexpression of *Fas2* recapitulated the anatomical phenotype of *miR-210*^*KO*^ flies, we decided to use the CRISPR-Cas 9 system to delete the *miR-210* binding site within the *Fas2* 3’UTR. As the seed region of miRNAs (positions 2-7), is critical for miRNA function [46], to minimize potential off-target effects, we managed to delete only 7 bps of the matching sequence on the 3’UTR (Figure 3B). We recovered one line, which is homozygous viable, and named the mutant: *Fas2*^*ΔmiR-210*^.

**Figure 3:**
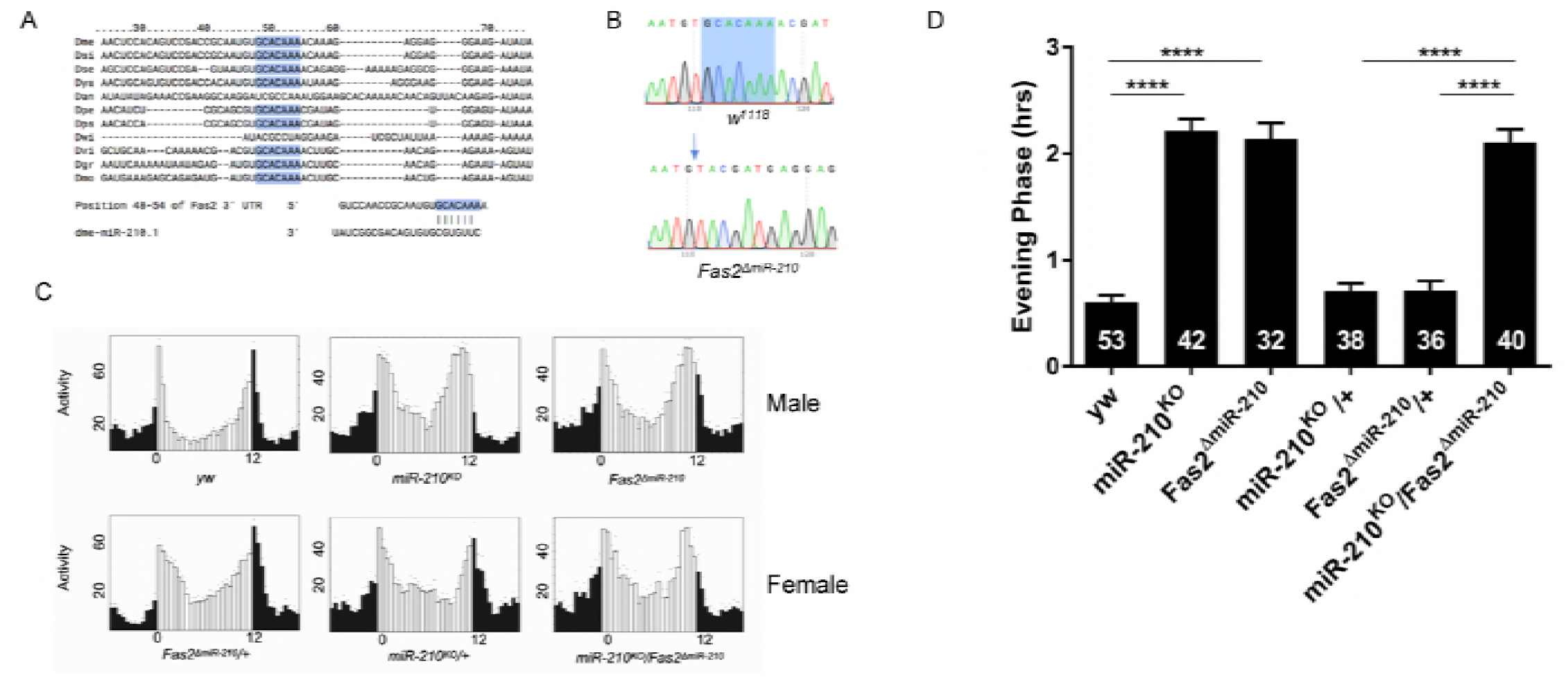
Depletion of putative *miR-210* binding sites within 3’UTR of *Fas2* phenocopies *miR-210*^*KO*^. (A) Predicted *miR-210* binding site in the 3’UTR of *Fas2* is highly conserved across *Drosophila* species. Blue letters indicate conserved sequences matching the 2-7 (seed) sequence of *miR-210*. (B) Confirmation of CRISPR/Cas9 mediated deletion of the 7bp from Fas2 3’UTR through Sanger sequencing. Blue arrow indicates the place that deletion started. (C) Representative eduction profiles of locomotor activity under 12:12 LD conditions. Since both *miR-210* and *Fas2* are located on the X chromosome, females were used in the low panel for genetic interactions. (D) Quantification of phase of evening peak. Error bars indicate Standard Error. **** = p<0.0001, determined by Student’s t test.

Next we tested the circadian locomotor behavior of the *Fas2*^*ΔmiR-210*^. There was no change in period under DD (Table 1). However, the *Fas2*^*ΔmiR-210*^ flies clearly advanced the phase of evening activity, similar to *miR-210*^*KO*^ flies (Figure 3C-3D). To determine whether Fas2 and *miR-210* function in the same pathway, we tested the potential genetic interactions. While both *miR-210*^*KO*^ and *Fas2*^*ΔmiR-210*^ are recessive mutations, the *miR-210*^*KO*^/*Fas2*^*ΔUTR*^ transheterzygous flies had a strikingly similar phase advance phenotype as single mutants (Figure 3C-3D). Taken together, *Fas2* genetically interacts with *miR-210*, and ablation of the *miR-210* binding sites within the 3’UTR of *Fas2* recapitulated the behavior phenotypes of *miR-210*^*KO*^ mutants. These results indicate that *Fas2* is a functional target of *miR-210*.

### *miR-210* regulates circadian rhythms through inhibition of Fas2

To identify the cellular mechanism of *miR-210* functions, we firstly characterized the *miR-210* expression pattern in the fly brain by using the *miR-210* knock-in Gal4 to drive GFP as a reporter. We observed GFP signals in the optic lobe, Hofbauer-Buchner eyelet (H-B eyelet), as well as several other brain regions, including the mushroom bodies and antennal lobe (Figure 4A). To our surprise, no GFP signal was detected in known circadian neurons as labeled by anti-PER antibody staining. Then we examined Fas2 expression using the commercial antibodies (1D4) from DSHB. As previously reported, in wild type flies Fas2 was strongly expressed in the mushroom bodies as well as the ellipsoid bodies (Figure 4B), [47]. Interestingly, we detected an intensive Fas2 signal in the optic lobe of *miR-210*^*KO*^ mutant flies, which disappeared when *miR-210* expression was rescued. In addition, elevated abundance of Fas2 was also observed in the optic lobe of *Fas2*^*ΔmiR-210*^ (Figure 4B-4D). Lastly, the increased Fas2 signal clearly overlapped with *miR-210* expression in the optic lobe (Figure 4E).

**Figure 4:**
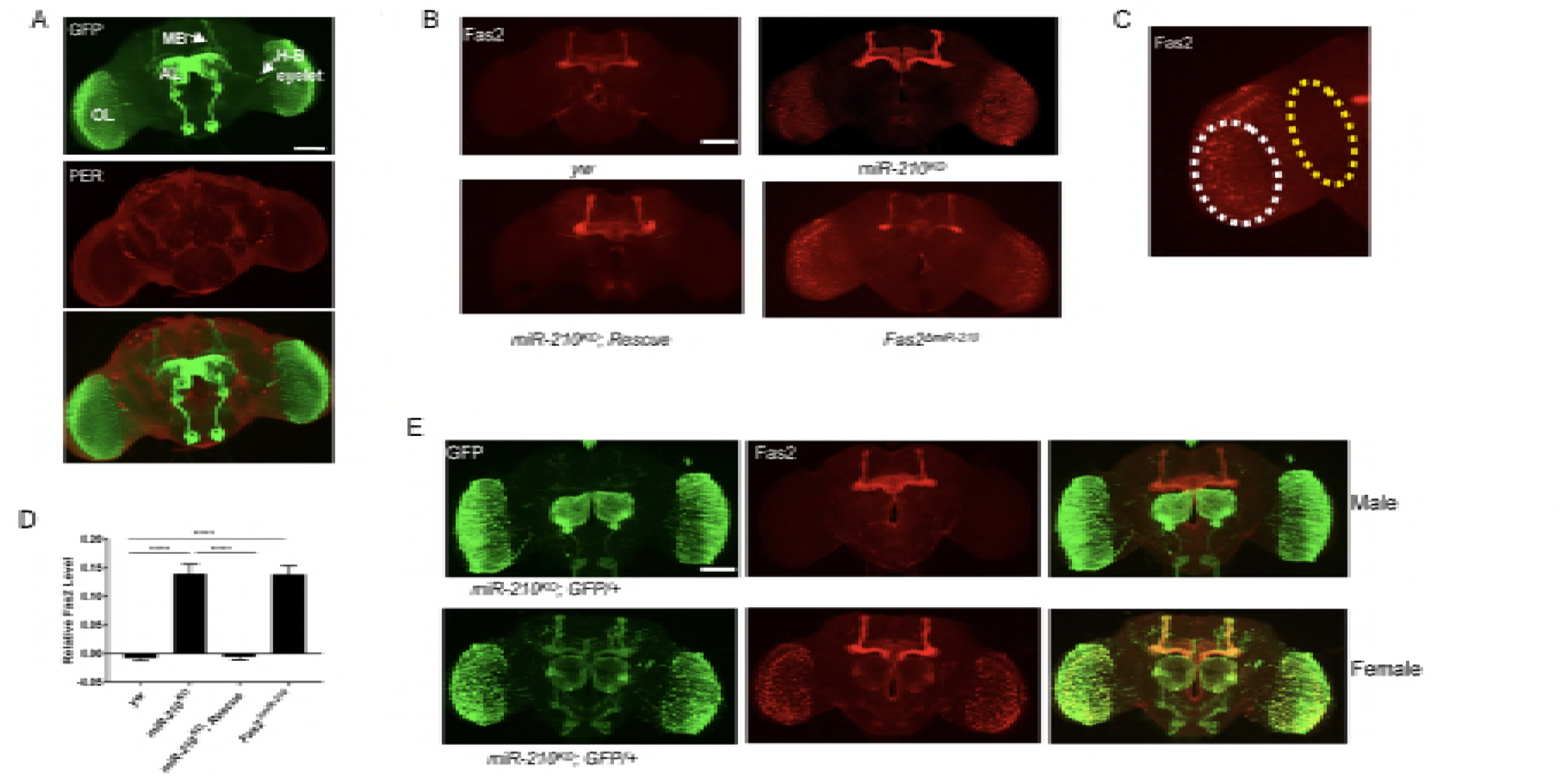
Fas2 abundance is elevated in the optic lobe of *miR-210*^*KO*^ and *Fas2*^*ΔmiR-210*^ flies. (A) Representative image showing *miR-210* expression profile in fly brain. Green signal represents *miR-210* expression, while red indicates PER. OL, AL, and MB represents optic lobe, antennal lobe and mushroom bodies, respectively. (B) Representative confocal images for Fas2 staining in fly brains. (C) Schematic representing area luminance analysis for quantification of relative Fas2 level in the optic lobe. Scale bar is 75 μm in A-C. (D) Quantification of Fas2 in the optic lobe per square μm. Error bars indicate Standard Error. **** = p<0.0001, determined by Student’s t test. (E) Loss of *miR-210* increases Fas2 levels in the cells where *miR-210* was expressed. Fly brains stained with anti-GFP (red) and Fas2 (green).

Although we did not detect miR-210 expression in circadian neurons with the *miR-210* knock-in Gal4, the single cell RT-PCR data from Chen and Rosbash indicated that *miR-210* has an oscillating expression in the sLNvs [48]. Thus, we first tested the requirement of *miR-210* in the sLNvs for normal circadian behavioral rhythms under LD, combining *miR-210* GAL4 and Pdf-GAL80. GAL80 is a repressor of GAL4 function, and Pdf-GAL80 has been efficiently used to inhibit gene expression in the PDF neurons [7, 49]. Surprisingly, we observed a clear rescue of evening phase when we excluded *miR-210* expression in the sLNvs, which suggests that *miR-210* is not necessary in PDF neurons for circadian behavior (Figure 5A). Next, we tested whether overexpression of Fas2 in the *miR-210-*expression cells would mimic the behavior phenotype of *miR-210*^*KO*^ mutants. We observed a weaker (∼1 hr) but significant advance of evening peak in Fas2 overexpression (Figure 5B-5C). Consistent with the *miR-210* data, Fas2 is not required in the PDF positive neurons (Figure 5).

**Figure 5:**
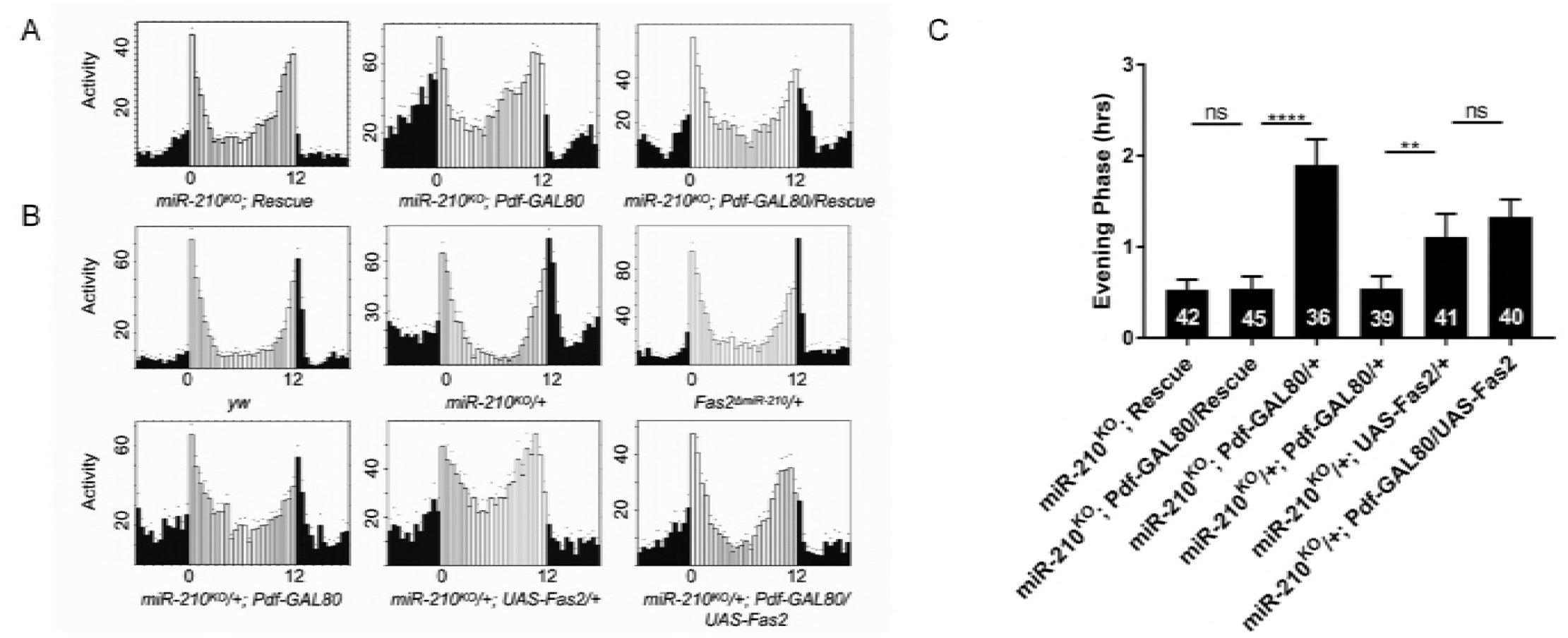
*miR-210* is required in the optic lobe for controlling the phase of evening behavior. (A-B) Representative eduction profiles of flies under 12:12 LD cycle. (A) *miR-210* expression in PDF negative cells is sufficient to rescue the advance in evening peaks. Overexpression of *Fas2* in PDF negative neurons advanced the phase of evening peaks. Female flies were used in this figure. (C) Quantification of evening phase. Error bars indicate Standard Error. ** = p<0.01, **** = p<0.0001 was determined by Student’s t test.

## Discussion

Emerging roles of miRNA in the control of different aspects of circadian rhythms have been recently uncovered [33]. Here, we screen *Drosophila* miRNA mutants and demonstrate that *miR-210* is critical for circadian locomotor rhythms. *miR-210* determines the proper phase of evening activity peak under entrainment. *miR-210* plays its role in circadian rhythms and axonal arborization via repression of *Fas2*.

*Drosophila* gradually increases its locomotor behavior and reaches its peak of evening activity at light off. The molecular mechanism underlying the phase control of evening peak is still largely unknown. Loss of *miR-210* advanced the evening phase to the same extent of *pdf*^*01*^ mutants. In fact, our genetic interaction data suggests that *miR-210* functions in the same signaling pathway as *pdf*. However, unlike the *pdf*^*01*^ mutants, loss of *miR-210* has no effect on morning anticipation or circadian rhythmicity. These results indicate that *miR-210* may specifically regulate one aspect of *pdf* signaling functions.

Circadian structural plasticity of sLNv dorsal projections has been recently identified [50, 51]. The biological function and molecular mechanisms underlying the structural plasticity however remain elusive. Here we identified that *miR-210*^*KO*^ mutants disrupted the axonal arborization rhythms of sLNv, which is likely through inhibition of the neural cell adhesion molecule Fas2. Deletion of the 7-bp seed region of *miR-210* binding sites within the 3’UTR of *Fas2* mimics the effect of *miR-210*^*KO*^ mutants on the sLNv arborization rhythm. However it seems unlikely that the phase advanced phenotype of *miR-210*^*KO*^ mutants is because of the defects in sLNv arborization rhythms. First, as far as we know, previous genetic manipulations that cause severe defects on sLNv structural plasticity have no effect on evening phase [50, 51]. Second, although overexpression or down regulation of Fas2 in the sLNv abolished the arborization rhythms, it does not affect the evening phase either [15]. So we conclude that these effects of *miR-210* on evening phase control and axonal structural plasticity maybe not functional related.

Our evidence that *Fas2* is a key target for *miR-210* in regulation of circadian rhythms is particularly strong. First, *miR-210*^*KO*^ mutants elevated the Fas2 abundance in the optic lobe, where *miR-210* is highly expressed. This is consistent with that miRNAs are usually negative regulators of target protein. In addition, *Fas2*^*ΔmiR-210*^ flies also mimic the increase of Fas2 level as observed in *miR-210*^*KO*^ flies. Second, *Fas2*^*ΔmiR-210*^ recapitulates the anatomical and circadian behavior phenotypes of loss of *miR-210*. Furthermore, genetic interaction data also indicates that *Fas2* and *miR-210* function in the same pathway.

In which cells are *miR-210* and *Fas2* required for the evening phase of activity? Transcriptome profiling from the Rosbash lab show that expressions of both *miR-210* and *Fas2* are enriched and oscillating in the PDF positive neurons [15]. Overexpressing *miR-210* in the PDF neurons causes flies arrhythmic or rhythmic with a prolonged circadian period (Table 1). However, using *miR-210* knock-in GAL4 or Fas2 antibodies, we were not able to detect obvious signals for *miR-210* or Fas2 in the PDF neurons. We cannot exclude the possibility that there is weak amount of *miR-210* or Fas2 protein expression, which is not detectable due to sensitivity of the techniques we used. We sought to answer this question by utilizing the well-used Pdf-GAL80 repressor. Indeed, restoring *miR-210* in its endogenous expressing cells except PDF neurons rescued the circadian behavior completely. This data indicates that expression of *miR-210* in the PDF neurons is not necessary for its circadian function. Consistent with this notion, overexpression of Fas2 in PDF negative but *miR-210* expressing cells causes a phase advance of evening peak. Since the *miR-210* knock-in driver has no detectable expression in the PDF neurons, potential leaky expression due to weak suppression of Pdf-GAL80 should not be an alternative explanation. The H-B eyelets interact with lLNvs and control the phase of evening peak activity [52, 53]. Interestingly, *miR-210* exhibits a strong signal of expression in the H-B eyelets (Figure 4A). It is possible that *miR-210*^*KO*^ mutants disrupt the interaction between H-B eyelets and lLNv, thus affect the evening activity.

Here we identify that *miR-210* regulates circadian locomotor rhythms and axonal structural plasticity of pacemaker neurons in *Drosophila*. A similar mechanism may exist in mammals. Interestingly, the glutamatergic synapses on the VIP neurons of mammalian master circadian pacemaker suprachiasmatic nucleus (SCN) also show circadian structural plasticity, which maybe important for light entrainment in mice. *miR-210* is a highly conserved miRNA from worms, flies, to humans. *In silico* miRNA target prediction with targetscan identified a highly conserved binding site of human *miR-210* in the 3’UTR of vertebrates *brain-derived neurotrophic factor* (BDNF). BDNF has well-established functions in axonal branching and synaptic plasticity [54, 55]. Thus, it is worthwhile to test that whether *miR-210* plays conserved functions in structural plasticity in mammals.

## Material and Methods

### Fly stocks

All the flies were raised on standard cornmeal/agar medium at 25 °C under 12:12 hour LD cycle. The following strains were used in this study: *w*^*1118*^, *yw, y w; tim-GAL4/CyO* [40], *y w; Pdf-GAL4/CyO [7*], *UAS-miR-210, UAS-Fas2, miR-210*^*KO*^ [43], *Fas2*^*ΔmiR-210*^*(generated by CRISPR-Cas9 in our lab).*

### Behavioral experiments and analysis

Most of the times, adult male flies (2-5 days old) were used to test locomotor activity rhythms. Since both miR-210 and Fas2 are both on the x chromosome, adult female flies were also used for locomotor rhythms, which were mentioned in the relative figures. Flies were entrained for 4 days LD cycle at 25 °C and 60% humidity, and released into constant darkness (DD) at 25 °C for at least 5 days. Temperature cycles was performed with 12h:12h 29C:21C cycling for 6d in complete darkness. Trikinetics *Drosophila* activity monitors were used to record locomotor activity in I36-LL Percival incubators. Activity rhythms were analyzed with Faasx software protocol [56]. Eduction profiles were generated with 3d of LD activity. FAAS-X software was used to analyze behavioral data [56]. Actograms were generated with a signal-processing toolbox for MATLAB. The morning anticipation amplitude was determined by assaying for the locomotor activity as described.

### Immunohistochemistry and quantification

Immunohistochemistry was performed with whole Drosophila brains. For staining involving ZT timepoints, Flies were entrained to LD for 3 d and dissected at Zeitgeber time (ZT) 0, 1, or 14. For CT stainings, flies were entrained to LD, released to DD, and dissected on the second day of DD at 6 CT intervals. Mouse anti-Fas2 (1:100), mouse anti-PDF (1:400), Rabbit anti-GFP 1:200, Rabbit anti-PER (1:1500) was used. All brains were imaged with a leica confocal microscope. Microscope laser settings were held constant for each experiment. ImageJ was used for quantification of PER, PDF and FAS 2 luminance. At least 5 brains were used for quantification. ImageJ was used in the analysis of axonal morphology (fasciculation) of sLNv dorsal termini by a modified Sholl’s analysis.

### Design of CRISPR mediated deletion and screening of Fas2 3’UTR deletions

*Fas2*^*ΔmiR-210*^ flies were generated by CRISPR mediated deletion by Rainbow Transgenic Flies. The guide RNA target sequence for CRISPR was GCAATGTGCACAAAACGATGAGG. The primers used for genomic *miR-210* f3: GCCAACAGGCAGCATCAAAC, and *miR-210* r3: CCAACTTAGTGTGCCAATCGATC. PCR amplification of the Fas2 3’ UTR region containing the *miR-210* motif was performed by using *miR-210* f3 and *miR-210* r3 primers. Sequencing of the PCR products was performed by using the *miR-210* f3 primer. Identification of homozygous *Fas2*^*ΔmiR-210*^ flies was confirmed by observing a 7bp deletion at the *miR-210* binding motif in the sequence chromatograph.

## Acknowledgements

We thank members of the Zhang lab for technical supports and discussions. We thank Dr. Patrick Emery and Dr. Junhai Han for carefully reading the manuscript and suggestions. We thank Dr. Michael Rosbash for the *UAS-Fas2* fly strains. *Pdf-Gal80* and *pdf*^*01*^ flies are generous gifts from Dr. Patrick Emery. We also thank Dr. Ralf Stanewsky for the anti-PER antibodies. We thank the Bloomington stock Center for various fly stocks, including *miR-210*^*KO*^, *UAS-miR-210, Rh1-Gal4, MB-Gal4*. We also thank the Developmental Studies Hybridoma Bank for PDF antibodies. Yong Zhang’s lab is supported by the National Institutes of Health COBRE Grant P20 GM103650.

## Author Contributions

Y.Z. formed the idea, supervised the project and designed the experiments. W.L., X.N., and Z.L. performed the experiments and analysis. Y.Z., W.L. wrote the manuscript.

## Supporting Information

**S1 Fig. *miR-210* genetically interacts with *pdf.*** (A) Representative eduction profiles of fly locomotor activity under 12:12 LD cycle. Black represents the dark phase, while white represents the light phase. Eduction is analyzed based on average of 3 days LD. (B) Quantification of evening phase of flies under LD. Number of flies tested is listed in each bar. (C) Representative eduction profiles under 16:8 LD cycle. (D) Quantification of evening phase of flies under 16:8 LD. Number of flies tested is listed in each bar. Error bars indicate Standard Error. ****= p<0.0001, determined by Student’s t test.

**S2 Fig. The molecular pacemaker is intact in the *miR-210*^*KO*^ mutants.** (A) Representative images of sLNvs in wild-type control and *miR-210*^*KO*^ mutants. Fly brains were dissected at six time points (circadian time, CT) during the second day of DD and stained with anti-PDF (green) and anti-PER (red) antibodies. (B) Quantification of PER staining in sLNv. No significant changes in PER level or cycling. Error bars indicate SEM. (C) Quantification of PDF levels in sLNv at CT1. Error bars indicate SEM. ns=no significance, determined by Student’s t test.

**S3 Fig. PER oscillation is not affected in the DN1 and LNds of *miR-210*^*KO*^ mutants**. (A-B) Representative images of DN1 (A) and LNd (B). Fly brains were dissected and stained same as Figure S2. (C-D) Quantification of PER staining in DN1 (C) and LNd (D). Error bars indicate SEM. ns=no significance, determined by Student’s t test.

**S1 Table. Locomotor activity of miR[KO] flis in DD.**

